# A ‘phenotypic hangover’ – the predictive adaptive response and multigenerational effects of altered nutrition on the transcriptome of *Drosophila melanogaster*

**DOI:** 10.1101/152330

**Authors:** Amy J. Osborne, Peter K. Dearden

## Abstract

The Developmental Origins of Health and Disease (DOHaD) hypothesis predicts that early-life environmental exposures can be detrimental to later-life health, and that mismatch between the pre- and postnatal environment may contribute to the growing non-communicable disease (NCD) epidemic. Within this is an increasingly recognised role for epigenetic mechanisms; epigenetic modifications can be influenced by, e.g., nutrition, and can alter gene expression in mothers and offspring. Currently, there are no whole-genome transcriptional studies of response to nutritional alteration. Thus, we sought to explore how nutrition affects the expression of genes involved in epigenetic processes in *Drosophila melanogaster.* We manipulated *Drosophila* food macronutrient composition at the F0 generation, mismatched F1 offspring back to a standard diet, and analysed the transcriptome of the F0 – F3 generations by RNA-sequencing. At F0, the altered (high protein, low carbohydrate, HPLC) diet increased expression of genes involved in epigenetic processes, with coordinated downregulation of genes involved in immunity, neurotransmission and neurodevelopment, oxidative stress and metabolism. Upon reversion to standard nutrition, mismatched F1 and F2 generations displayed multigenerational inheritance of altered gene expression. By the F3 generation, gene expression had reverted to F0 (matched) levels. These nutritionally-induced gene expression changes demonstrate that dietary alteration can upregulate epigenetic genes, which may influence the expression of genes with broad biological functions. Further, the multigenerational inheritance of the gene expression changes in F1 and F2 mismatched generations suggests a predictive adaptive response (PAR) to maternal nutrition. Our findings may help to understand the interaction between maternal diet and future offspring health, and have direct implications for the current NCD epidemic.

## Background

Exposure to aberrant or harmful environments during development and early life can be detrimental to later life health. From these observations is derived the Developmental Origins of Health and Disease (DOHaD) hypothesis (Barker, 2007, Gluckman and Hanson, 2004a, Gluckman et al., 2010, Bruce and Hanson, 2010), which seeks to explain why the period from conception to birth and the first few years of life is critical for determining life-long susceptibility to non-communicable diseases (NCDs). Many NCD phenotypes are thought to be caused by developmental perturbations that are a consequence of altered epigenetic marks (Gluckman and Hanson, 2004a, Heindel, 2006), induced by environmental exposure during critical periods of development (Skinner et al., 2013). Alteration to the epigenome regulates gene expression through DNA methylation, histone and chromatin modifications (Callinan and Feinberg, 2006, Peaston and Whitelaw, 2006), providing plasticity to the genome. Consequently, phenotypes under epigenetic regulation provide a pathway though which the genome can interact with the environment (Holliday and Pugh, 1975). If the epigenetic modifications occur at a time during which they are able to affect the germline, such modifications may also influence development of the offspring (Skinner et al., 2013).

The interaction between the environment and the epigenome and the resulting phenotypic adaptations, coupled with the growing NCD epidemic, led to the predictive adaptive response (PAR) hypothesis (Gluckman and Hanson, 2004b). This hypothesis states that nature of the predictive adaptive response (PAR) is determined by the degree of mismatch between the pre- and postnatal environments. This mismatch results from the information that the fetus receives on environmental conditions while *in utero,* to which it will respond adaptively by programming its biology to expect that environment If the actual postnatal environment matches the prenatal prediction, then the PARs are appropriate and disease risk is low; if they do not match then the PAR is inappropriate, and disease risk is increased. For instance, obesity has a distinct epigenetic profile. This pattern could be established in early life as a response to the maternal, fetal and/or early postnatal environment, and later-life nutritional mismatch could mean that the individual has been programmed inappropriately, leading to an increased risk of obesity and associated diseases later in life (Cordero et al., 2015). This implies that the epigenetic hallmarks of early life exposures may be able to be maintained or ‘stored’ in such a way as to produce long-lasting effects (Lee, 2015).

Along with stress, drugs and environmental toxicants, one of the main factors that can cause epigenetic perturbation is nutrition; evidence suggests that early-life nutrition can not only affect the long-term health of the individual, but also of their offspring (Tarry-Adkins and Ozanne, 2011, Langley-Evans, 2015, Lillycrop and Burdge, 2015) potentially through epigenetic mechanisms (Aiken and Ozanne, 2013, Haggarty, 2013, Vickers, 2014a, Vickers, 2014b, Vaiserman, 2014). Consistent with the DOHaD hypothesis, strong links exist between both maternal and early life nutrition and cardiovascular disease (Barker, 1997), diabetes and obesity (Uauy et al., 2011) along with asthma and allergy, autoimmune disease, cancer and mental health (Gluckman et al., 2011, Barouki et al., 2012, Hanson and Gluckman, 2015, Balbus et al., 2013). Inappropriate maternal nutrition has been linked to incorrect epigenetic ‘priming’ during fetal or postnatal life (Burdge et al., 2007b, Brudasca and Cucuianu, 2016); in particular, diets high in carbohydrate content (sugar) have been shown to program metabolic status and diabetes (Buescher et al., 2013, Musselman et al., 2011).

Many nutrition-related phenotypes have been attributed to changes in epigenetic processes and a classic example of this is that of methyl supplementation, which influences coat colour in agouti mice (Waterland and Jirtle, 2003). Nutrition influences are separated into direct effects and also indirect (offspring) effects. Direct effects can be exemplified by high-fat diets inducing obesity and metabolic syndrome, due to the methylation pattern of particular genes and promoters such as leptin (Milagro et al., 2009) and PPARγ (Fujiki et al., 2009), and the differential DNA methylation detected in metabolic syndrome in both humans and rodents (Luttmer et al., 2013, Sánchez et al., 2015). Indirect effects on offspring metabolic phenotype can occur via maternal diet. For example, high fat diets in utero are known to affect offspring epigenetic patterns and methylation status of particular genes, for example, adiponectin and leptin genes (Masuyama et al., 2016), while a high fat diet during pregnancy and lactation can induce epigenetic modifications and differential expression of the μ-opioid receptor, and corresponding hypomethylation of the promoter regions of the gene, in mouse offspring (Vucetic et al., 2010). Additionally, maternal protein restriction can cause hypomethylation of particular genes involved in metabolic processes in fetus and offspring (Lillycrop et al., 2007, Burdge et al., 2007a, van Straten et al., 2010, Burdge et al., 2004) and can also affect methylation in the developing placenta (Reamon-Buettner et al., 2014). Such epigenetic perturbation is not just limited to fetal and early life; the postnatal period is also susceptible to the epigenetic effects of nutrition. For example, hypermethylation of the pro-opipmelanocortin promoter occurs in over-fed rats (Plagemann et al., 2009), and postnatal folic acid supplementation can lead to hypermethylation of PPARα (Burdge et al., 2009). This sensitivity of the epigenome to the effects of the environment (nutrition) also extends into adulthood, where epigenetic changes have been observed in response to nutritional changes (Christman et al., 1993, Waterland et al., 2006, Hoile et al., 2013).

In addition to metabolic genes, altered nutrition appears to have broader genomic consequences. For instance, nutrition can also alter markers of inflammation and oxidative stress (Jacometo et al., 2015); a protein restricted diet in pregnancy leads to an increased susceptibility to oxidative stress in offspring (Langley et al., 1994), while a high carbohydrate diet increases the oxidative stress response (Gregersen et al., 2012). A high fat, high carbohydrate meal can induce oxidative and inflammatory stress as reflected by increased reactive oxygen species (ROS) generation in both normal weight (Aljada et al., 2004) and in obese people (Patel et al., 2007), suggesting that oxidative stress and inflammation are major mechanisms involved in metabolic disorders associated with obesity (Fernández-Sánchez et al., 2011) and can also induce epigenetic changes (Sánchez et al., 2015). Indeed, the stress response is well established to be under epigenetic control in humans (Cencioni et al., 2013). In plants, DNA and histone modifications control gene expression when a plant is under environmental stress (Chinnusamy and Zhu, 2009). In terms of applicability to health, we know that low doses of reactive oxygen species, from calorie restricted or high carbohydrate diets, promote health and life span (Ristow and Schmeisser, 2011). Thus considering the above, along with the PAR hypothesis, it is likely that such biological responses to nutrition reflect the idea that induced epigenetic changes that underpin physiological change, and aid in the adaptation of an individual, and potentially its offspring, to an adverse environment (Gluckman et al., 2005).

Nutritional deficiency may also lead to the development of psychopathological behaviour such as antisocial, violent and criminal behaviour (Neugebauer et al., 1999). This may in part be a consequence of the capacity of nutritional deficiency to alter brain development (Liu and Raine, 2006), possibly through epigenetic factors, which may lead to changes in brain structure and function (Liu et al., 2015). Supporting this is the suggestion that macronutrient deficiency can cause changes in epigenetic regulation, which can lead to impaired brain development, signalling molecule imbalance, neurotoxicity and differences in neurotransmission (Liu et al., 2015) which all contribute to psychopathological outcomes. For example, studies into participants from the Dutch Hunger Winter cohort found correlations between prenatal famine exposure and schizophrenia (Susser and Lin, 1992, Susser et al., 1996), along with persistent epigenetic changes at, e.g. the *IGF1* gene (Heijmans et al., 2008).

Thus, considering that gene-specific studies of altered nutrition have demonstrated broad and diverse genetic and epigenetic consequences, it is pertinent to apply this concept to the whole genome. Nutrition is commonly investigated as an environmental factor that is expected to influence the epigenetic landscape, and there are several examples in the literature of response to altered nutrition being ‘inherited’ multigenerationally (Burdge et al., 2007b, Jirtle and Skinner, 2007, Carter et al., 2004, Golding, 2004) and transgenerationally (F3 and beyond, (Waterland and Jirtle, 2003)). As such, altered gene expression, via epigenetic marks in response to nutrition, coupled with the PAR hypothesis, could be the key to understanding the prevalence of obesity and metabolic syndrome. Here we explore the PAR hypothesis and the ability of nutrition to affect gene expression at a whole-genome level by manipulating the diet of the fruitfly, *Drosophila melanogaster,* to investigate the extent to which gene expression is changed by differing levels of macronutrients. Previous research has shown that a high-sugar maternal diet can alter the body composition of larval *Drosophila* offspring for at least two generations (Buescher et al., 2013), as well as demonstrating that nutrition is able to influence traits relative to metabolic syndrome, longevity and the immune response (Musselman et al., 2011, Morgan, 2012, Roussou et al., 2016). Considering such traits and responses are often under epigenetic control, we predict that dietary manipulation will have broad consequences for the expression of genes involved in epigenetic processes.

## Methods

### Fly husbandry

*Drosophila melanogaster* stocks used in this study were wild-type Canton-S flies from the Bloomington Drosophila Stock Center at Indiana University. *Drosophila* were cultured in a dedicated invertebrate laboratory using standard techniques. Briefly, flies were maintained in laboratory incubators at 25C in a P Selecta HOTCOLD-C incubator. Larvae were reared on either a standard low protein high carbohydrate (LPHC, standard laboratory fly food) or a high protein low carbohydrate (HPLC) diet. These differential diets consisted of standard brewer’s yeast (Health2000, NZ), sugar (New Zealand Sugar Company, Auckland, New Zealand) and cornmeal (Health2000, NZ), in varying ratios (Table 1). Agar (A7002, Sigma-Aldrich, MO, USA), propionic acid (Thermo Fisher, New Zealand, AJA693) and Nipagin (47889, Sigma-Aldrich, MO, USA, 10% w/v in 100% ethanol) were added in equal amounts. Gross energy (KJ/g) of both the LPHC and HPLC food types was determined by bomb calorimetry, and total protein content (%) was calculated using the total combustion method (Table 1) by the Institute of Food, Nutrition and Human Health at Massey University, Palmerston North, New Zealand.

**Table 1.** Fly diet components and content information

| **Component** | **High Protein** | **Low protein** |
| --- | --- | --- |
| Agar (g) | 9 | 9 |
| Cornmeal (g) | 66.7 | 66.7 |
| Sugar (g) | 31.24 | 46.7 |
| Yeast (g) | 148.76 | 16.7 |
| Propionic acid (ml) | 6.6 | 6.6 |
| Nipagen (ml) | 5 | 5 |
| Water (L) | 1 | 1 |
| Protein (%) | 8 | 5.3 |
| Gross energy (KJ/g) | 0.9 | 2.1 |
| Yeast:sugar ratio | 01:00.2 | 01:02.8 |

### Nutrition experiments

*Drosophila* were manipulated under anaesthesia (CO_2_) and initially raised on LPHC (low protein, high carbohydrate, standard fly) food. To enable mating, 50 female flies were segregated within 4 hours of eclosion, and incubated with 10-15 male flies, on LPHC food, for 24 hours. Female flies were separated and incubated on either LPHC or HPLC diet. Female flies laid eggs in their specified food and the F1 offspring from the HPLC diet were either maintained on HPLC, or mismatched onto LPHC diet (Figure 1). The further offspring then remained on those matched or mismatched diets, relative to the F0 generation. Thus the biological mismatch relates to flies from a HPLC (non-standard) dietary background that are mismatched onto a LPHC background.

**Figure 1.**
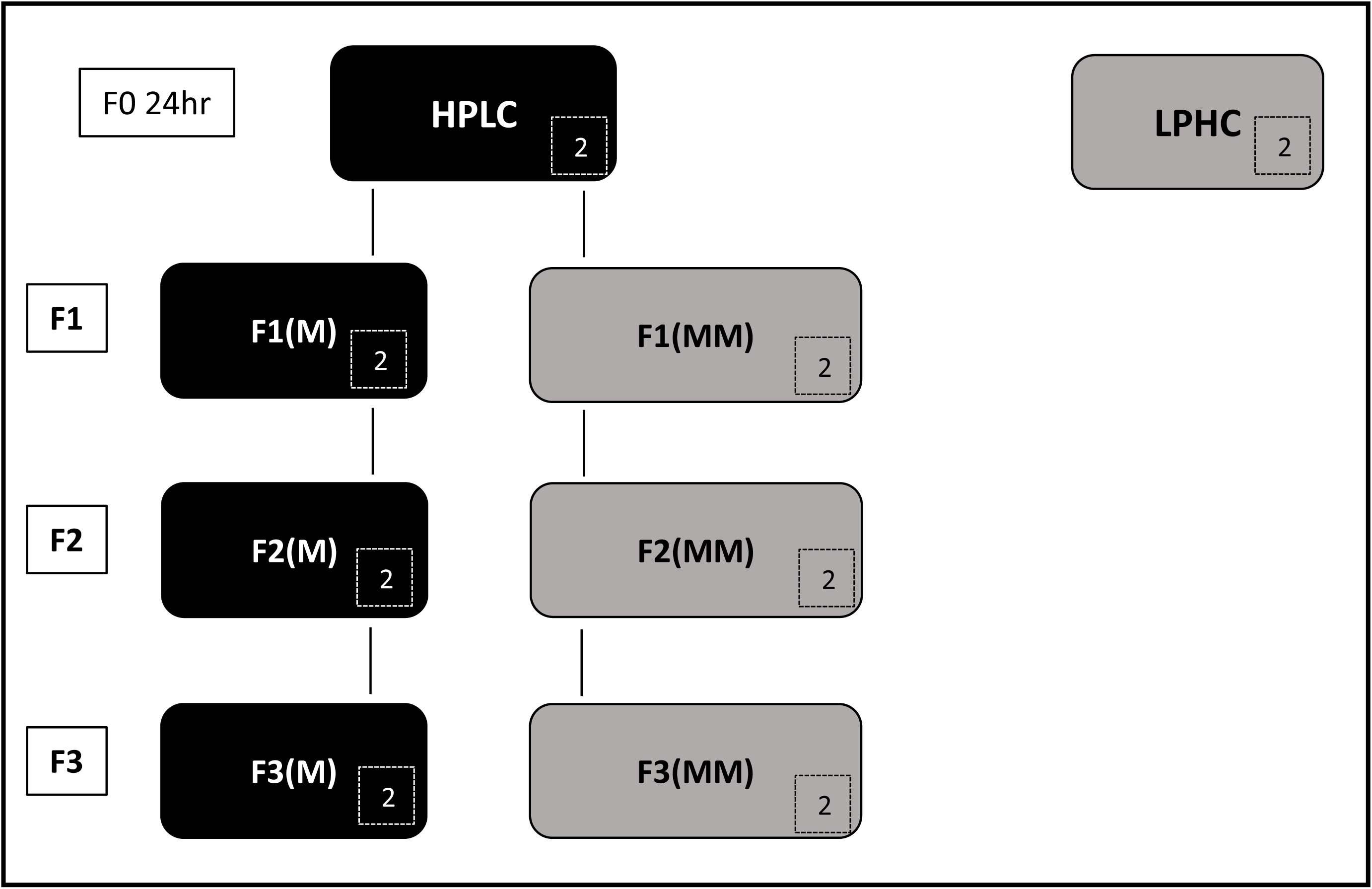
Fly diet experiments. LPHC, low protein, high carbohydrate (standard) diet; HPLC, high protein low carbohydrate diet; F1(M), F2(M) and F3(M), flies maintained on LPHC diet for three generations; Fl(MM), F2(MM) and F3(MM), flies that were raised on HPLC in the F0 generation, and mismatched back to LPHC at éclosion in the F1 generation, and were maintained for two more generations on the mismatched (LPHC) diet. Boxed numbers denote the number of replicates used for each condition for transcriptomic experiments.

At each generation, RNA was extracted from female *Drosophila* using a modified kit protocol. Firstly, 10 female *Drosophila* were ground in 300ml of Trizol reagent (Invitrogen) that had been cooled on dry ice, with a disposable, sterile pestle until homogenised. A further 200μl of Trizol was added and the homogenised tissue was incubated in Trizol reagent for 5 min. 200 μl of Chloroform was added and the solution was vigorously mixed via vortex for 15 seconds. This solution was incubated on ice for 5 min and centrifuged at 12,000 g at 4°C for 10 min to separate the aqueous (containing RNA) and organic phases. The aqueous solution was removed and added to an equal volume of 70% ethanol and mixed via pipetting. The Qiagen RNeasy Kit (Qiagen, Hilden, Germany) was then used to purify the RNA, according to the manufacturer’s instructions. RNA was stored at -80°C until needed.

### Transcriptomic experiments

Two F0 generation replicates from each of the LPHC and HPLC diets and two replicates from the F1, F2 and F3 generations with matched and mismatched diets were prepared (boxed numbers, Figure 1), resulting in 16 total RNA samples submitted to the Otago Genomics and Bioinformatics Facility at the University of Otago (Dunedin, New Zealand) under contract to the New Zealand Genomics Limited for library construction and sequencing. The libraries were prepared using TruSeq stranded mRNA sample preparation kit according to the manufacturer’s protocol (Illumina). All libraries were normalised, pooled and pair-end sequenced on 2 lanes of high-output flowcell HiSeq 2500, V3 chemistry (Illumina), generating 100-bp reads. Libraries had an average insert size of ~208bp.

Transcriptomic output was analysed in CLC Genomics Workbench Version 8.5.1. Reads were aligned to the *Drosophila melanogaster* reference genome (BDGP6) as implemented in CLC Genomics Workbench, and differential gene expression (EDGE test) was calculated between samples using an absolute fold change value of >1.5, and an FDR-corrected p-value of <0.001. Transcriptomic data was validated by Nanostring: samples were submitted to the Otago Genomics and Bioinformatics Facility at the University of Otago (Dunedin, New Zealand) under contract to the New Zealand Genomics Limited for nCounter Custom Gene Expression assays (Nanostring). 100ng total RNA in 5uL total volume was processed using the standard nCounter XT Total RNA protocol. The Total RNA and the CodeSet were combined with hybridisation buffer and incubated at 65C for 20 hours. The Codeset consists of Reporter and Capture probes that hybridise the target sequence of interest, forming a tripartite complex. Hybridised samples were then processed in batches of 12 using the High Sensitivity Protocol (3 hours). Raw data was exported and QC-checked using Nanostring’s nSolver data analysis tool (www.nanostring.com). As per the Nanostring CodeSet design criteria, 25 candidate genes for validation were chosen, including two housekeeping genes incorporated (Mnf and Rpl32, Table 2). Raw data was normalised to the geometric mean of both the positive controls (included in the hybridisation steps) and the nominated housekeeping genes. Normalised Nanostring data was compared to transcriptomic data and the Pearson’s correlation coefficient was calculated in R (Team, 2008).

**Table 2.**
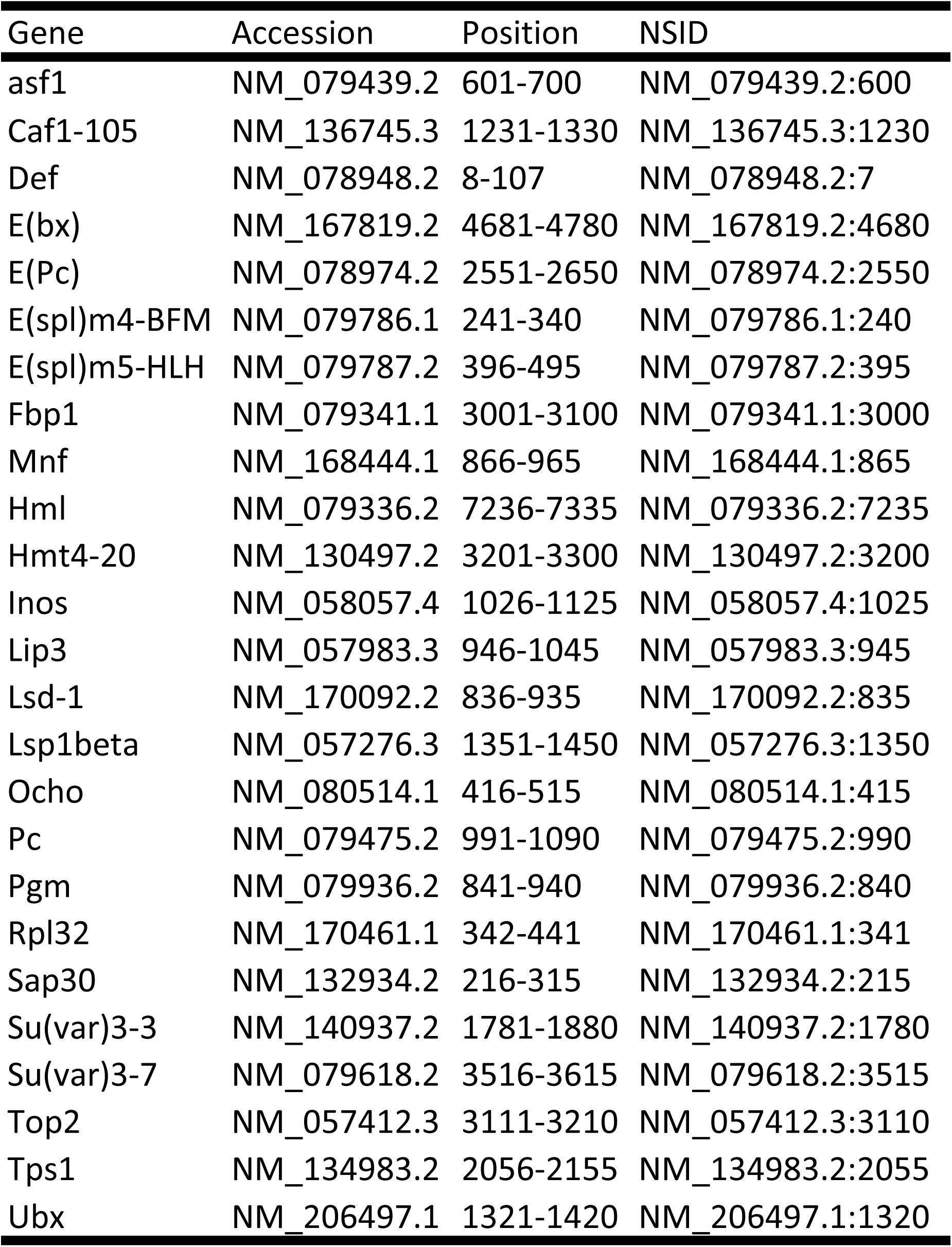
Codeset design for Nanostring. Gene, gene symbol based on FlyBase nomenclature; NSID, Nanostring internal identifier; Position, region in the target mRNA being probed

### Gene ontology (GO) analyses

Functional annotation clustering (FAC) was undertaken in the Database for Annotation, Visualization and Integration of Discovery (DAVID) v6.8 (Huang et al., 2009b, Huang et al., 2009a) with the following categories: COG_ONTOLOGY, GOTERM_BP_DIRECT, GOTERM_CC_DIRECT, GOTERM_MF_DIRECT, KEGG_PATHWAY and INTERPRO. Up- and down-regulated genes from the F0 generation were submitted separately to DAVID for FAC, and analysed against a background of genes that were expressed and detectable in this dataset, to identify GO terms that were significantly enriched between the HPLC and LPHC F0 generation.

### Statistical analyses

Significant differences in mean gene expression between different dietary conditions and generations was calculated via ANOVA and Tukey’s posthoc testing, as implemented in R (Team, 2008).

## Results

### RNAseq data

Summary statistics for transcriptomic work is shown in Supplementary file 1. Briefly, each sample yielded between 3164 Mbases and 4286 Mbases (average 4001 Mbases), with an average number of reads of 32,006,648 reads (range 25,310,758 – 34,288,584 reads). The mean quality score was an average of 36 PF (range 35.4 – 35.72 PF).

### Differential gene expression

Of 17,490 genes annotated in the Drosophila genome and contained within the CLC reference database, 12,424 were expressed and detected in these transcriptomic experiments. Of these 12,424 genes, 2946 were differentially expressed (1074 [8.6%] down-regulated and 1872 [15.1%] up-regulated (Supplementary file 2), with an absolute fold change of >1.5, and an FDR-corrected p-value of <0.001, as determined by EDGE test as implemented in CLC.

### Gene ontology

DAVID uses a clustering approach to reduce redundancy; GO terms that are similar, are clustered together. Each term within the cluster is given a p-value, and the cluster itself is given an enrichment score (the geometric mean in –log scale of the individual GO term p-values). An enrichment score of 1.3 is equivalent to a non-log p-value of 0.05.

Functional annotation clustering of genes that are up-regulated in HPLC vs. LPHC indicated that the dataset was highly enriched for genes that are involved in epigenetic processes such as chromatin binding (ES 10.64), DNA replication (enrichment score, ES, 4.64), chromatin regulation (ES 1.62) and histone binding and phosphorylation (ES 1.60) (Table 3). Conversely, clustering of genes that are downregulated in HPLC vs. LPHC indicated that the dataset was highly enriched for genes that are involved in immunity (ES 11.41), fatty acid metabolism (ES 3.49), neurotransmission (ES 3.35) cellular metabolic processes (ES 2.49) and oxidative stress pathways (ES 2.19, Table 4).

**Table 3.**
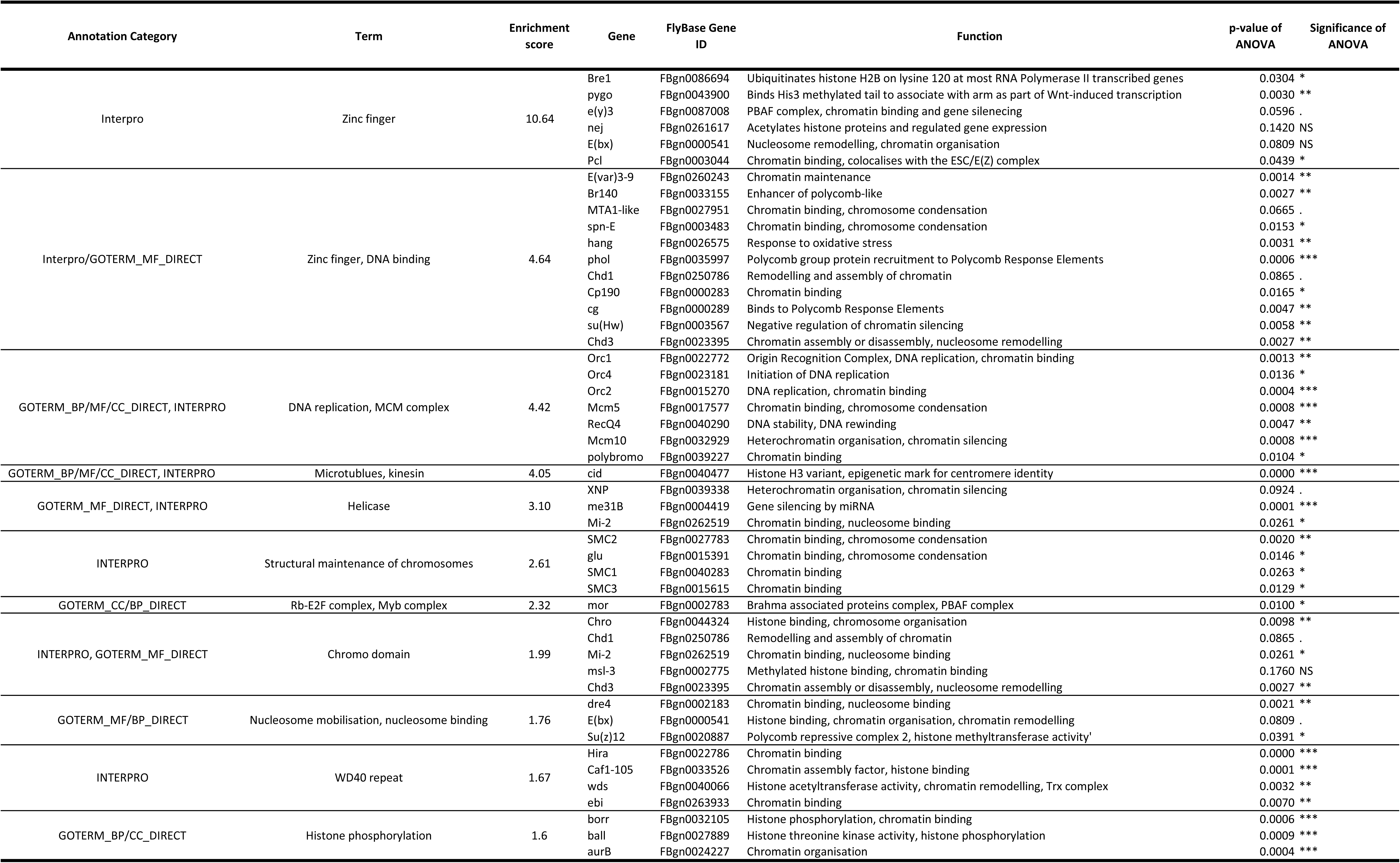
Functional annotation clustering (FAC) as performed in DAVID. Genes that were significantly up-regulated in these data in HPLC compared to LPHC were compared to a background list of genes that were expressed and detected in these data. Analyses of variance were carried out on expression between all samples in this study (F0-F3 matched and mismatched) with p-values and statistical significances listed.

### Validation

Based on the genes that were upregulated in this study, we selected a panel for 20 genes for transcriptomic data validation. All transcriptomic samples were validated by Nanostring, per-sample r value of 0.81-0.95, whole dataset correlation r of 0.86 (Supplementary file 3).

### Multigenerational gene expression of genes of interest

To determine whether the pattern of gene expression that we identify as being altered in response to diet is maintained across generations (F0-F3, matched [M] and mismatched [MM]) when the HPLC diet is removed, we analysed the multigenerational expression levels of particular epigenetic genes of interest that were identified as being upregulated in HPLC. From the lists of clustered genes generated by DAVID, the same pattern of expression was observed for every gene, which showed the intermediate-level maintenance of the upregulation of the genes in the F1 and F2 generation, followed by a reversion to F0 (matched) gene expression levels by the F3 generation. Figure 2 displays a selection of indicative graphs which display this effect, with ANOVA significance data listed in Table 3 (described in Supplementary file 4) and significant pairwise comparisons as determined by Tukey’s posthoc testing (Supplementary file 5) indicated by solid and dashed lines. There is no significant difference between the expression of epigenetic genes when comparing the F0 LPHC diet and the F3MM flies, despite an intermediate and significant difference between F1 and F2 flies mismatched onto LPHC diets.

**Figure 2.**
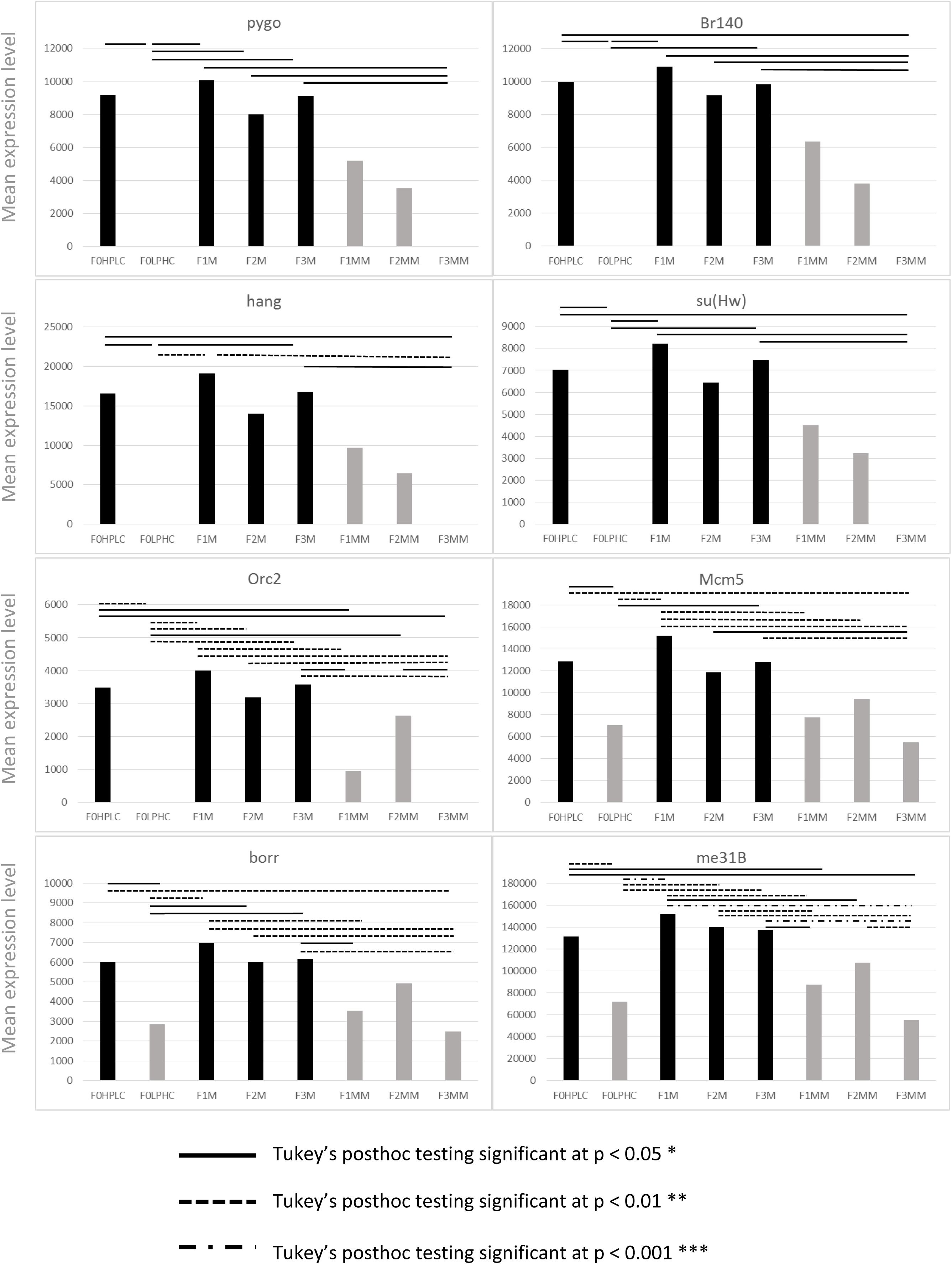
Indicative graphs of gene expression of genes up-regulated in the F0 generation, with significances as determined by ANOVA, between F0-F3 generations on matched and mismatched diets. Pairwise comparisons by Tukey’s posthoc testing indicated by solid and dashed lines, as described in the key. Y axis denotes mean expression level from transcriptomic experiments, X axis denotes the dietary condition as per Figure 1. Note differing Y axis scales. Gene names are stated as per their FlyBase gene symbol IDs.

When we consider genes that were significantly downregulated between the F0 flies, we see the same pattern of gene expression, in the opposite direction. Choosing to look at genes that are clustered using DAVID (Table 4) we see that genes that are downregulated in response to diet, remain at a low level in the F1 and F2 mismatched cohorts (F1MM and F2MM) but by F3, their gene expression has regained the same level as the LPHC (F0) generation, despite being lower in the F1 and F2 generations (Figure 3 and Supplementary files 4 and 6). This effect is genome-wide, and applies to every gene tested from the lists generated by DAVID.

**Figure 3.**
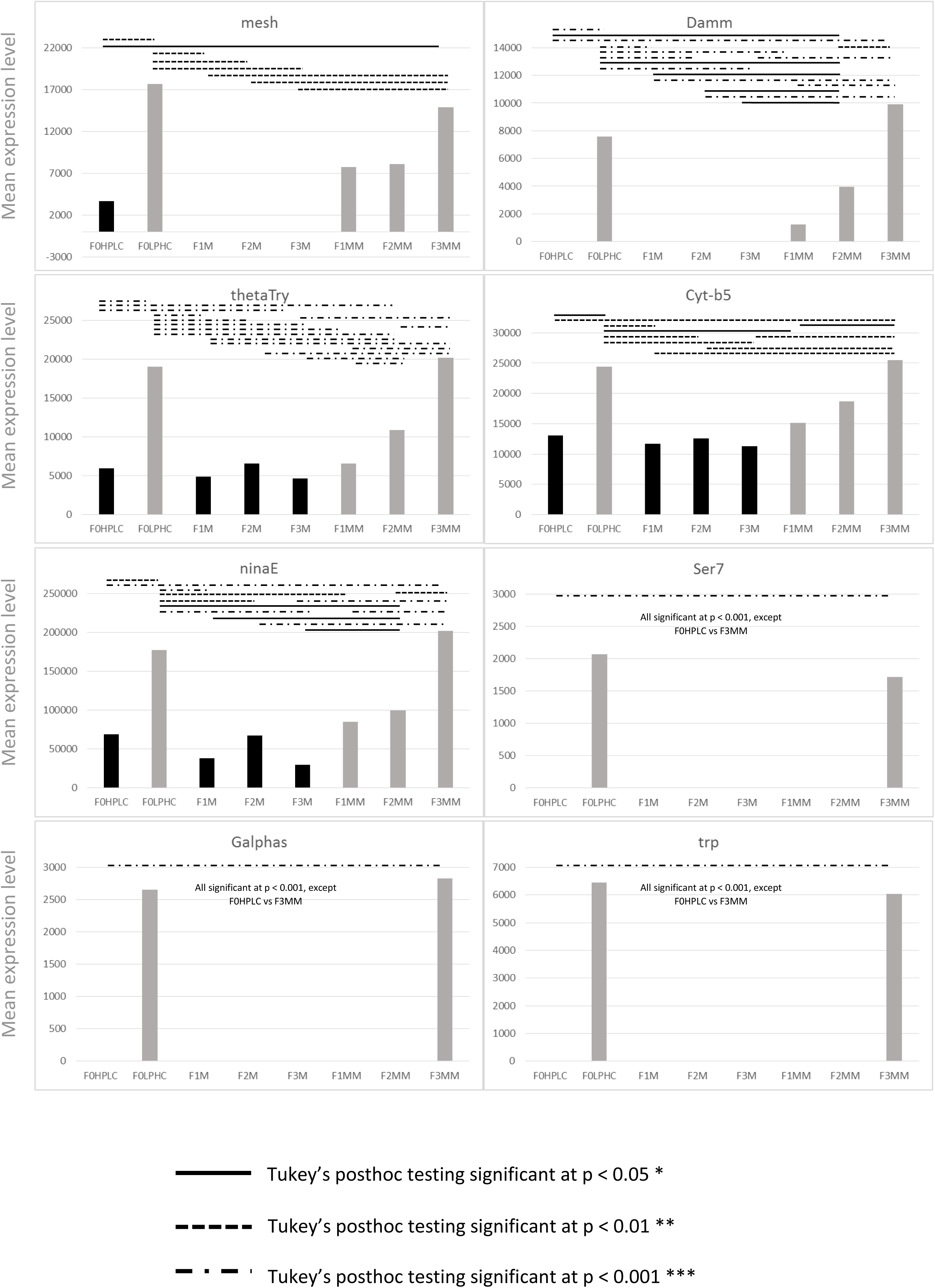
Indicative graphs of gene expression of genes down-regulated in the F0 generation, with significances as determined by ANOVA, between F0-F3 generations on matched and mismatched diets. Pairwise comparisons by Tukey’s posthoc testing indicated by solid and dashed lines, as described in the key. Y axis denotes mean expression level from transcriptomic experiments, X axis denotes the dietary condition as per Figure 1. Note differing Y axis scales. Gene names are stated as per their FlyBase gene symbol IDs.

**Table 4.**
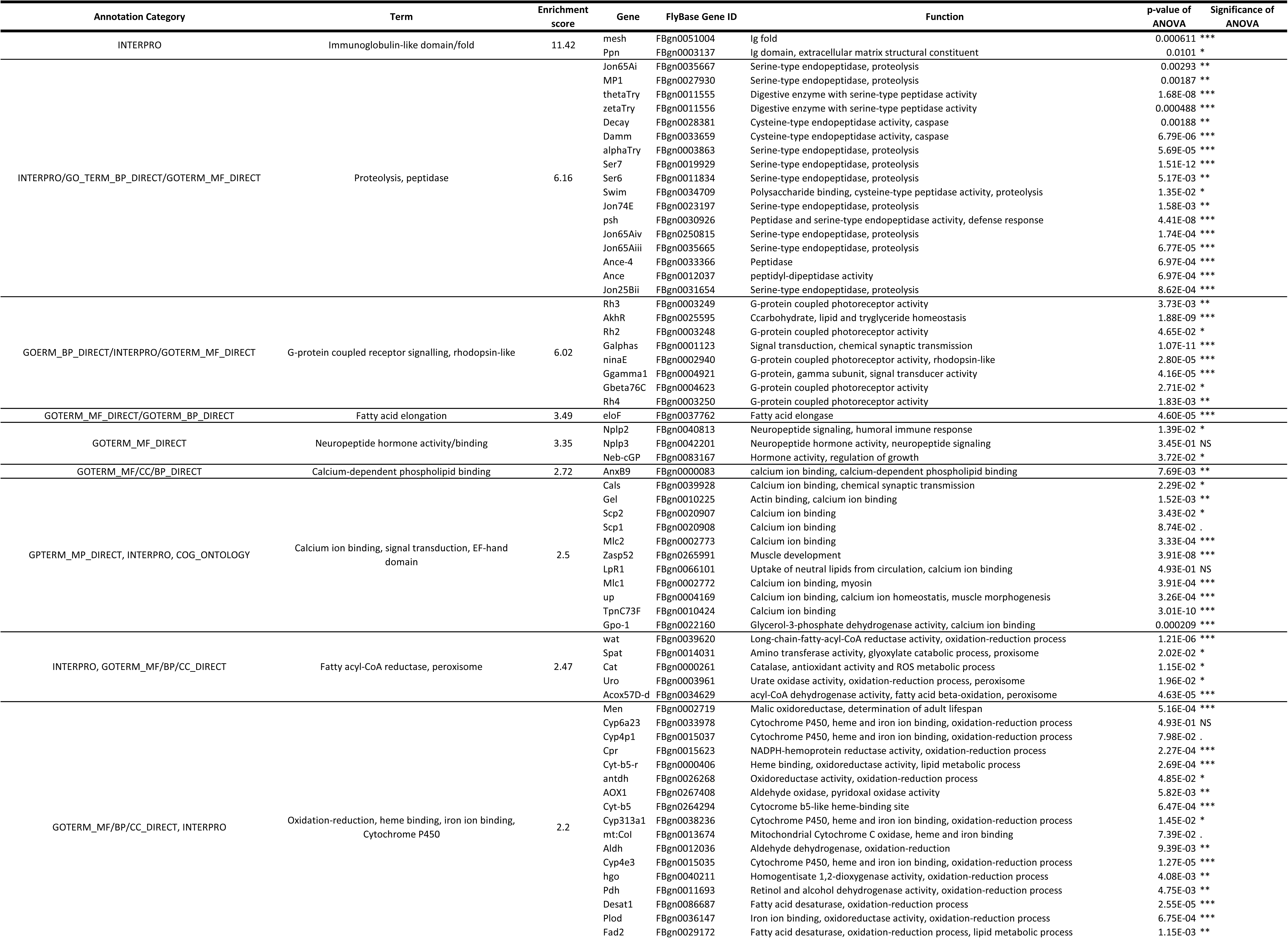
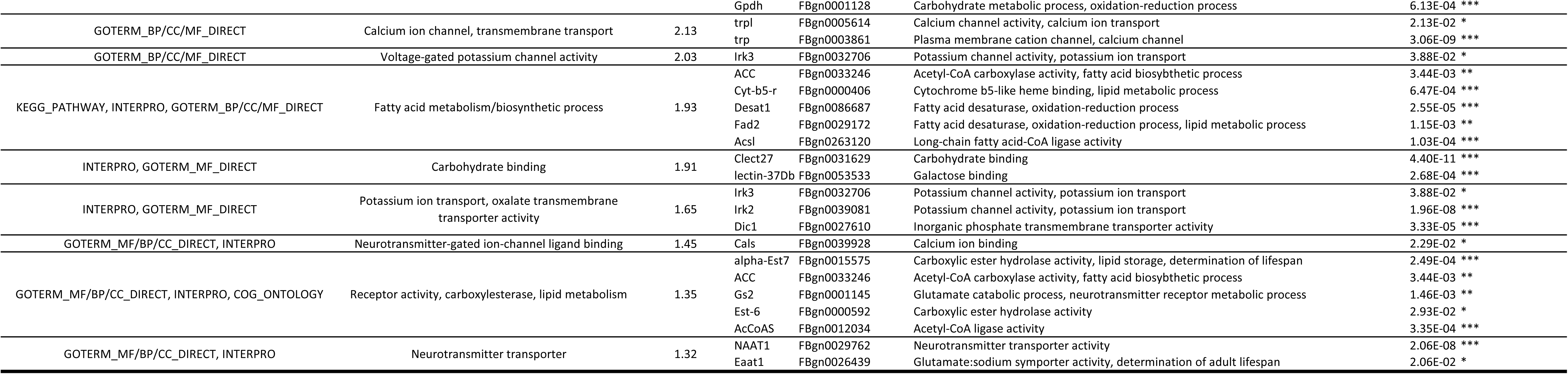
Functional annotation clustering (FAC) as performed in DAVID. Genes that were significantly down-regulated in these data in HPLC compared to LPHC were compared to a background list of genes that were expressed and detected in these data. Analyses of variance were carried out on expression between all samples in this study (F0-F3 matched and mismatched) with p-values and statistical significances listed.

## Discussion

The primary goal of this study was to investigate the effect of nutrition on the expression of genes involved in epigenetic processes. We have demonstrated that a high protein, low carbohydrate (HPLC) diet results in genome-wide upregulation of epigenetic genes in the F0 generation, compared to that observed in a standard *Drosophila* low protein, high carbohydrate (LPHC) diet; this effect was so strong that the overwhelming majority of genes that were upregulated in response to diet were involved in epigenetic processes, with very few other classes of genes categorised as significantly upregulated. Classes of genes that were downregulated in response to the HPLC diet, versus LPHC, were broader in scope, and included genes involved in the immune response, cell signalling, oxidative stress, carbohydrate and fatty acid metabolism, and neurotransmission. Thus, this study shows that altering a LPHC diet to a HPLC results in genome-wide upregulation of genes involved in epigenetic processes, with a coordinated down regulation of genes with broad physiological functions. The coordinated nature of the gene expression data we observed imply, firstly, that genes involved in the processes of neurotransmission, oxidative stress, metabolism and immunity appear to be under epigenetic control, and secondly, that this epigenetic control, and altered biological response, is influenced by nutrition.

In response to the genomic and epigenomic changes observed in the F0 generation, we further questioned if dietary alteration resulted in developmental programming of the biological response to diet; specifically, whether the changes induced by the HPLC diet in the F0 generation persisted beyond F0, upon removal of the HPLC diet. The genomic changes induced in the F0 generation persisted at intermediate levels in mismatched F1 and F2 generations, in the absence of the HPLC diet. By the F3 generation, gene expression in the mismatched flies had reverted back to the level observed in the F0 matched generation. We hypothesise, firstly, that this multigenerational inheritance of gene expression, followed by a reversion to matched F0 levels by the time the F3 generation is reached, indicates moderate epigenetic programming in the form of a predictive adaptive response (PAR). We further hypothesise that the ‘correction’ of the genomic changes induced by dietary alteration after three generations implies that dietary reversion to match the F0 generation may be able to correct an altered genomic landscape, effectively rescuing an aberrant nutrition-induced phenotype.

The environment is able to interact with genes through epigenetic mechanisms (Holliday and Pugh, 1975) particularly during development (Gluckman and Hanson, 2004a, Gluckman et al., 2010, Bruce and Hanson, 2010); it is during development, when DNA is replicating at an accelerated rate, that the genome is most sensitive to environmental influences that could induce epigenetic alteration, which may influence gene expression and the development of the animal. Crucially, this could also lead to the alteration of the epigenome of the germ cells (Skinner et al., 2013). Any permanent alteration to the germ cell epigenome (Guerrero-Bosagna et al., 2010) may then be transmitted through the germ line, with adverse phenotypic consequences for offspring (Barker, 2007, Skinner, 2011). For example, adult-onset diseases can be induced through embryonic exposure to environmental toxins, primarily endocrine disruptors (Anway et al., 2006, Newbold et al., 2006, Heindel, 2006, Anway and Skinner, 2006). Thus, if epigenetic modifications do become permanent, these modifications can be inherited by future generations and impact disease susceptibility (Anway et al., 2005, Jirtle and Skinner, 2007).

A large number of studies report transgenerational inheritance in a range of eukaryotes (reviewed in (Jablonka and Raz, 2009)). Many of these studies, particularly in mammals, report inheritance of the acquired trait over two or three generations. Concordant with work by Jirtle and Skinner (2007), we agree that these effects should not be defined as truly transgenerational, because, mechanistically, exposure of an F0 gestating female to an environmental stimulus (nutrition, toxicants, stress) also exposes the F1 embryo (Figure 1, (Osborne et al., 2013)). Further, for species that develop *in utero,* parental exposure also exposes the germ cells that will form the F2 generation. Thus traits present in the F2 generation should be considered as multigenerational, rather than transgenerational, as they could have been induced by direct environmental exposure through the fetus and the germ line. To reflect this, some studies searching for evidence of true transgenerational inheritance are declining to assay the F1 generation entirely (Xia et al., 2016) due to the fact that transmission to the F1 generation can be indicative of both parental effects and programming (Xia and de Belle, 2016). Thus, here we describe our findings from the F1 and F2 generations as multigenerational inheritance, since the gene expression changes induced by the environment in the parental generation revert to F0 levels after the F2 generation.

The current NCD epidemic is etiologically very complex but it is thought to be mediated, in part, by developmental aberrations arising from the inheritance of altered epigenetic marks (Gluckman and Hanson, 2004a, Heindel, 2006). Many metabolic phenotypes and gene expression differences are linked to differential epigenetic marks that are nutritionally-induced. For example, a protein-restricted diet during pregnancy causes hypomethylation of the hepatic PPARα (peroxisome proliferator-activated receptor alpha) and glucocorticoid receptor (GR) genes in rats, and promotes the same hypomethylation in the F1 and F2 offspring of F0 rats fed a protein-restricted diet during pregnancy, despite the nutritional challenge being only in the F0 generation (Burdge et al., 2007b). Others have reported evidence of embryonic environmental exposure influencing the phenotype of the F1 generation (Carter et al., 2004, Golding and Team, 2004, Kang et al., 2002, Klip et al., 2002, Portha, 2005, Tsui and Wang, 2004), as well as, specifically, maternal nutrition exerting effects on the F1 phenotype (Carter et al., 2004, Golding and Team, 2004). Such research strongly implies that epigenetic effects could be the key to understanding the current epidemic of overweight and obese, and associated metabolic syndromes, particularly if nutrition in the F0 generation can induce a PAR to nutrition, as we hypothesise is occurring here. Interestingly, comprehensive studies using animal models that investigated the effect of both protein restricted and energy-rich diets during pregnancy on the phenotype of the offspring showed that offspring born to dams fed these different diets exhibited persistent metabolic changes, similar to those observed in human metabolic disease such as obesity, insulin resistance and hypertension (Lillycrop and Burdge, 2015), indicating an element of developmental programming and a possible PAR. These findings imply that both famine (protein restriction) and energy-rich diets, when mismatched back to adequate nutrition, are similarly detrimental to the metabolic health of offspring, and that it is possibly the mismatch itself between inadequate nutrition and proper nutrition which is leading to metabolic disease phenotypes. This highlights the fact that epigenetic mechanisms play a highly complex role in obesity and metabolic pathways (Xue and Ideraabdullah, 2016, Lillycrop and Burdge, 2011, Lillycrop and Burdge, 2015, Li et al., 2010).

In addition to the striking expression level changes observed in genes involved in epigenetic processes, one major source of change that we observed in these data are genes involved in oxidative stress. Malnutrition or excess of particular nutrients can cause oxidative damage (Fang et al., 2002). For example, hyperglycaemia, which is an excess of sugar in the blood and one of the hallmarks of diabetes, is linked to a diet that is rich in carbohydrates and fat (Gupta et al., 2013, Yang et al., 2012). A build-up of sugar can lead to tissue damage, and this can be maintained because of metabolic memory (Sánchez et al., 2015), which itself may induce epigenetic changes and altered gene expression (Ceriello, 2010, El-Osta et al., 2008, Sánchez et al., 2015). Given that increased carbohydrate intake can induce oxidative stress as reflected by increased reactive oxygen species (ROS) generation (Aljada et al., 2004, Patel et al., 2007), our results are consistent with the observation that high carbohydrate diets are implicated in increased oxidative and metabolic stress (e.g. (Langley et al., 1994), and that the genome may be responding adaptively to dietary stressors. We know that oxidative stress responses are often under epigenetic control (Cencioni et al., 2013), and also, that maternal nutritional deficiency in pregnancy can lead to altered methylation and increased oxidative DNA damage in the brains of adult offspring (Langie et al., 2013), which, as well as being directly influenced by nutrition, may predispose to neurological disorders in later life.

Consistent with, and leading on from this observation, these data display a decrease in the expression of genes involved in neurotransmission and neurodevelopment when exposed to a HPLC diet. There is strong evidence linking oxidative stress to neurodegeneration and neurodegenerative disease, such as Alzheimer’s disease (Tabaton and Tamagno, 2007). In addition, it is also clear that an increase the production of ROS, induced by environmental factors, can increase risk of a multitude of neurodegenerative diseases (Migliore and Coppedè, 2009). Thus, it stands to reason that, in these data, nutrition may be impacting on the production of ROS and the expression of genes involved in ROS pathways and those involved in neurotransmission and neurodegeneration.

In addition to the DOHaD hypothesis, there are also free radical early life theories, which link environmental agents (e.g. diet, heavy metals) with perturbations of gene regulation and expression (in, for example, the *APP* gene), and the onset of, e.g., Alzheimer’s disease (Lahiri et al., 2007). Free radical early life theories also link the necessity for oxygen in histone demethylase action to epigenetic processes in development (Hitchler and Domann, 2007). These theories are supported by the observation that nutrition during pregnancy can induce epigenetic changes that result in altered nervous system development (McGowan et al., 2008) and also offspring cerebral function (Gallagher et al., 2005). Further, nutrient availability during the pre and postnatal periods can lead to long-lasting changes in neuron development (Niculescu et al., 2004), as well as influence the development of psychopathological behaviour (Neugebauer et al., 1999). This is because nutritional deficit may lead to altered brain development (Liu and Raine, 2006), possibly via epigenetic factors that can lead to changes in brain structure and function (Liu et al., 2015). Given that micronutrient availability can heavily influence neurotransmission, due to the fact that the function of the brain is inherently related to its metabolism of nutrients (Liu et al., 2015) in the form of vitamins and minerals that function as co-enzymes in neurotransmission and neurotransmitter metabolism, our gene expression data are supportive of these linkages.

Thus, through our data, we hypothesise that the genome-wide changes we observe in genes involved in epigenetic pathways could be responsible for the gene expression changes in other, broad, biological process seen in response to diet. The intermediate maintenance of these gene expression changes, even when the HPLC diet is removed, suggests a PAR to diet; the biology of the mismatched flies is programmed to expect a certain type of diet, and is responding adaptively, with altered gene expression in the absence of HPLC, albeit at a slightly lower level. The complete reversion of this in the F3 generation suggests an element of phenotypic rescue, implying that altered nutrition did not affect the germline, and that the gene expression changes are not fixed transgenerationally, and thus may have the capacity to be corrected over time.

To date, there have been no genome-wide assessments of the effect of nutrition on total gene expression. This study contributes to our understanding of the myriad ways in which nutrition can influence gene expression, phenotypes and future health outcomes, with relevance to the DOHaD hypothesis and the current NCD epidemic. While our study demonstrates multigenerational inheritance of gene expression values, rather than transgenerational, it is worth noting that the phenotypic effects of gene expression changes, rather than the gene expression itself, can persist and show multigenerational, and potentially transgenerational, inheritance. For example, a low protein diet given to *Drosophila* can increase H3K27me3 through upregulation of the E(z) protein. Interestingly, while the upregulation of the (E)z protein was not detected in the F2 generation, the associated increase in methylation H3K27me3 was in fact detected in the F2 generation (Xia et al., 2016) and the coordinated effect on longevity was also present through to the F2 generation. This suggests that while the gene expression and protein level is not inherited *per se,* the effects and/or functions of those genes possibly could be. It is possible that a phenomenon such as this may be present in these data; a permissive state may be achieved, whereby we might not detect gene expression changes inherited to F3 and beyond, but we may see associated genomic conformational or phenotypic changes in F3 and beyond. Further functional studies based on dietary manipulation are required to confirm this. In particular, it will be pertinent to prove causality between an epigenetic alteration and a change in regulation of genes involved in the traits we observe. To do so, we suggest a combination of phenotypic measures such as assessing lifespan, oxidative stress resistance, and immunity, as well as exploiting mutant *Drosophila* strains for genes of interest, to assess the effect of epigenetic alterations on downstream gene expression and associated phenotypes.

The mechanisms by which environmental perturbations affect the phenotype of the individual in the next generation are vital to understand. While hints of transgenerational epigenetic inheritance in more natural situations exist (e.g. (Jablonka and Raz, 2009)), more work is required to determine if an evolved mechanism exists for this transmission and its significance for the health, development, and evolution.

## Funding

This work was supported by a Gravida grant (MP04) to P.K.D.

## Acknowledgements

Stocks obtained from the Bloomington Drosophila Stock Center (NIH P40OD018537) were used in this study. This work was supported by a Gravida grant (MP04) to P.K.D. Under New Zealand law, *Drosophila* research is waived from requiring ethical approval, therefore none were needed for this study. *Drosophila melanogaster* stocks were imported under the HSNO Approval number GMC001092. All *Drosophila* work was carried out in accordance with the regulations of the HSNO Act 1996, as described in Morgan, 2012. We are grateful to Professor Sir Peter Gluckman for his input into, and discussions around, this subject. AJO would like to thank Professor Martin Kennedy for hosting the latter stages of this work.

## Data availability

All data supporting this work is included in the supplementary material.

## Supplementary Files

Supplementary file 1

Supplementary file 1.xlsx

Summary statistics from RNA sequencing experiments.

All experiments were carried out on the Illumina platform. The libraries were prepared using TruSeq stranded mRNA sample preparation kit according to the manufacturer’s protocol (Illumina). All libraries were normalised, pooled and pair-end sequenced on 2 lanes of high-output flowcell HiSeq 2500, V3 chemistry (Illumina), generating 100-bp reads. Libraries had an average insert size of ~208bp.

Supplementary file 2

Supplementary file 2.xlsx

Drosophila genes that are significantly differentially expressed between HPLC and LPHC F0 generation.

12,424 Drosophila genes were expressed and detected in these transcriptomic experiments. Of these 12,424 genes, 2946 were differentially expressed (1074 [8.6%] down-regulated and 1872 [15.1%] up-regulated), with an absolute fold change of >1.5, and an FDR-corrected p-value of <0.001, as determined by EDGE test as implemented in CLC.

Supplementary file 3

Supplementary file 3.xlsx

Transcriptomic data validated by Nanostring:

100ng total RNA in 5uL total volume was processed using the standard nCounter XT Total RNA protocol. The Codeset consists of Reporter and Capture probes that hybridise the target sequence of interest, forming a tripartite complex. Hybridised samples were processed in batches of 12 using the High Sensitivity Protocol (3 hours). Raw data was exported and QC-checked using Nanostring’s nSolver data analysis tool (www.nanostring.com). As per the Nanostring CodeSet design criteria, 25 candidate genes for validation were chosen, with two housekeeping genes incorporated within this amount (Mnf and Rpl32). Raw data was normalised to the geometric mean of both the positive controls (included in the hybridisation steps) and the nominated housekeeping genes. Normalised Nanostring data (expression level) was compared to transcriptomic data (mean expression level) and the Pearson’s correlation coefficient was calculated in R

Supplementary file 4

Supplementary file 4.xlsx

Analyses of variance (ANOVA) of F0-F3 matched and mismatched dietary conditions, up and down regulated genes.

ANOVAs of genes that have been identified through FAC as being significantly up or downregulated in the FO generation (in HPLC, vs. LPHC). Df = degrees of freedom, Sum sq = sums of squares, Mean sq = mean squares, F value = ANOVA F value, Pr(>F) = p-value of significance of F, SIgnifcance = significance indicator. Gene name = FlyBase gene symbol.

Supplementary file 5

Supplementary file 5.xlsx

Pairwise comparisons/Tu key’s posthoc testing of significant ANOVAs of genes that were signficantly upregulated in the F0 generation.

These data indicate the significant differences between the means of the different nutritional states (matched/mismatched) and generations, diff = difference, lwr,upr = confidence intervals, p adj = p value adjusted for multiple testing.

Supplementary file 6

Supplementary file 6.xlsx

Pairwise com parisons/Tu key’s posthoc testing of significant ANOVAs of genes that were signficantly downregulated in the F0 generation.

These data indicate the significant differences between the means of the different nutritional states (matched/mismatched) and generations, diff = difference, lwr,upr = confidence intervals, p adj = p value adjusted for multiple testing.

